# Genome-wide characterisation of pathogenicity-related proteins in *Mycena citricolor,* the causal agent of the American Leaf Spot in coffee

**DOI:** 10.1101/2023.12.30.573698

**Authors:** N. L. Larke-Mejía, N. Arciniegas, F. Di Palma, C. A. Angel C, J. J. De Vega

**Affiliations:** Earlham Institute, Norwich Research Park, Norwich, NR4 7UZ, UK; Plant Pathology Discipline, National Coffee Research Center (Cenicafé), National Federation of Colombian Coffee Growers (FNC), Planalto km. 4, Vía Antigua Chinchiná-Manizales, Manizales, Caldas, Colombia; School of Biological Sciences, University of East Anglia, Norwich, NR4 7TJ, UK

**Keywords:** *Mycena citricolor*, Agaricales, American leaf spot, hemibiotroph, long-read assembly, genome annotation, secretome, virulence factors

## Abstract

*Mycena citricolor* is a fungus that causes the American Leaf Spot (ALS) disease in multiple hosts, including coffee and avocado. This hemibiotroph penetrates the plant through damage induced by oxalic acid. This can cause 20-90% crop losses in coffee depending on the environmental and production conditions. *M. citricolor* is the only known pathogenic species in the *Mycena* genus, a large group of saprophytic mushrooms. Comparing the saprophytic and pathogenic genomes can allow us to identify genetic machinery associated with the pathogen’s genome-wide functional acquisitions to cause disease.

To identify pathogenicity-related genes in *M. citricolor*, we analysed protein family copy-number variation, secretome prediction, and homology to known virulence factors in two *M. citricolor* assemblies, including a newly assembled and annotated long-read genome. We found that the pathogenic *M. citricolor* had a higher proportion of secreted genes expanded in copy-number, and expanded gene copies homologous to known virulence factors than the saprophytic *Mycena*. We shortlisted over 300 candidate genes in each *M*. *citricolor* assembly. Focusing on genes strongly regulated during plant interaction, we found over 100 candidates, primarily from multiple copies (up to 4-3 times) of 42 well-known virulence factors (e.g. MFS1, CUTA, NoxA/B, OLE1, NorA), plus a few clade-specific uncharacterised genes.

*M*. *citricolor* transition to a pathogenic lifestyle reflected genome-wide functional changes. *M*. *citricolor* seems to primarily depend on well-known virulence factors in large copy numbers, suggesting the molecular plant-interaction processes involved are like those of better-studied fungi. Hypothetically, the development of ALS resistance could mirror studied responses to these virulence factors.

## Introduction

The American Leaf Spot (ALS; **Figure 1**) is a plant disease firstly spotted on Colombian coffee plants in 1876 (Buller 1934; Carvajal 1939; Uribe-Arango 1946; Wang 1988; Granados-Montero *et al*. 2020). The disease was soon later described in other crops and countries around the Americas (Buller 1934; Carvajal 1939; Wang 1988; Granados-Montero *et al*. 2020). The ALS is caused by the phytopathogenic fungi *Mycena citricolor* (Berk. & M.A. Curtis) Sacc. (Syn: *Omphalia flavida* Maubl. & Rangel) (Wang 1999; Wang and Avelino 1999), a Mycenaceae: Agaricales. *M. citricolor* is an A1 pest of quarantine concern in Europe (EPPO 2018). *M. citricolor* is a plurivorous pathogen with a broad spectrum of hosts, including fruit, forest, and ornamental crops (Ploetz 2007; Granados-Montero *et al*. 2017). The ALS can significantly affect coffee and avocado production (Wang 1999). In coffee plantations, the ALS periodically causes between 20% to 90% crop losses in Central and South American countries. The fungus can infect all coffee plant tissues (especially leaves, branch tips, and fruits) at all physiological stages and ages (**Figure 1**). However, the ALS is not endemic to all coffee production zones. It is mostly restricted to humid and shaded regions with frequent rainfall and cloudiness throughout the year, and moderate temperature, where crop management and local topography also play an important role (CABI 1989; Avelino *et al*. 2007; Rivillas-Osorio and Castro-Toro 2011).

**Figure 1.**
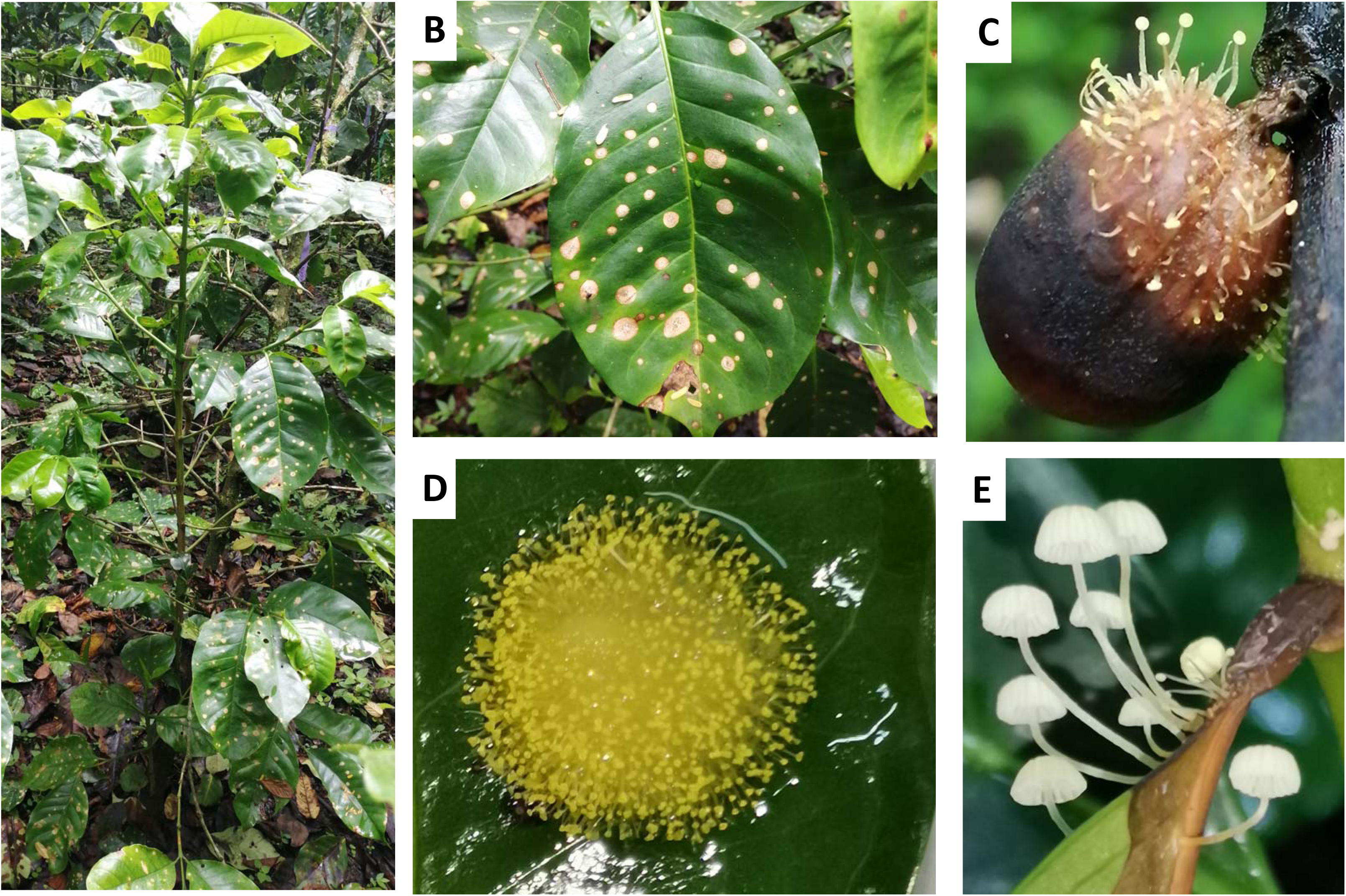
Images illustrating visible symptoms of the American Leaf Spot (ALS) on *Coffea arabica* (coffee plant) and various life stages of *Mycena citricolor*. A) Depicts a coffee plant exhibiting symptoms of ALS. B) Shows necrotic lesions, attributed to oxalic acid production by *M. citricolor* gemmae, on the leaf tissue of an ALS-affected coffee plant. C) Displays bonnet-shaped gemmae of *Mycena citricolor* growing on necrotised coffee bean fruit. D) Exhibits gemmae on coffee plant leaf cultivated in a laboratory setting. E) Represents basidiocarps, a feature of the sexual stage of *M. citricolor*.

The ALS develops when high humidity and water on the plant surfaces reactivate previous lesions and stimulate gemmae production and dispersion (**Figure 1C** and **1D**). Extreme weather events, such as the “La Niña” ENSO climatic pattern, which is associated with a prolonged rainy season in the central Andes and northern Brazil, favour severe epidemics from the ALS (Avelino *et al*. 2015). Higher weather variability associated with climate change is leading to an increased frequency of extreme “La Niña” events (Cai *et al*. 2015), affecting the incidence, severity and significant epidemics of the ALS in these regions. Currently, there are no coffee varieties resistant to ALS. Therefore, management strategies for this disease involve early inspection for symptoms, adjusting production systems (e.g., managing water and relative humidity levels, increasing solar radiation, airflow, and temperature in the crop), and periodically applying systemic fungicides (Angel *et al*. 2018). ALS epidemics are characterised by multiple infectious cycles occurring in a single season. These epidemics can also persist over several years, as the inoculum from one growing season carries over to the next (Angel *et al*. 2018).

*M. citricolor* has been described as a facultative hemibiotroph with a short biotrophic phase and a long necrotrophic phase. It is the only described plant pathogenic species in the *Mycena* genus. This genus includes widespread saprophytic litter and wood-decomposing bonnet mushrooms (Maas Geesteranus 1980), and several *Mycena* species evidenced endophytic association with plant seedlings with *in vitro* assays (Thoen *et al*. 2020). There is a thin line between pathogenicity and endophytism in some fungi, and many plant pathogenic fungi show a variety of interactions with their host plants at different stages (De Silva 2017; Salvatore 2020; DeMers 2022). Plant immune responses and the need for a more efficient mode of nutrient acquisition are proposed triggers for these lifestyle transitions (Andrew *et al*. 2012; Kabbage *et al*. 2015). Moreover, several fungal lineages have displayed evidence of multiple transitions between endophytic and pathogenic lifestyles, underscoring the dynamic nature of these interactions (Delaye *et al*. 2013; De Silva 2017).

The anamorph or asexual stage in *M*. *citricolor* is the most infective and common inoculum source in the field (García 2012). This phase comprises asexual mycelium which produces pin-shaped gemmiferous bodies formed by a gemma (tip) on top of a thin pedicel (**Figure 1C** and **1D**). The gemmae adhere to healthy plant tissue and secrete oxalic acid, initiating the degradation of epidermal cells, leading to the formation of necrotic lesions (**Figure 1B**) varying in colour from brown to dark purple (Rao and Tewari 1987). Oxalic acid, a pathogenicity factor in other plurivorous pathosystems, such as *Sclerotinia sclerotiorum* (Cessna *et al*. 2000b) and *Clarireedia jacksoni* (Townsend et al. 2020), plays a significant role in *M. citricolor*’s pathogenic process. Subsequently, *M*. *citricolor*’s new mycelia later develops on the necrotic tissue to generate new gemmae, which decapitate to cause secondary ALS infections. ALS lesions eventually coalesce resulting in premature leaf or fruit drop, necrosis on tips or branches, retarded plant growth, and in some aggressive cases, it can cause plant death (Buller 1934; Carvajal B 1939; Castaño A 1951; Angel *et al*. 2018). The sexual stage is a basidiocarp (sporophore) with lamella (**Figure 1E**), characterized by a small pale-yellow umbrella-shaped body, which produces rounded basidiospores (2n). The specific role of basidiospores in the infection process remains unknown (Sequeira 1954; Salas and Hancock 1972).

*M. citricolor* and other members of the large mycenoid clade have been studied using genomic approaches due to the independent evolution of bioluminescence within the clade (Kotlobay *et al*. 2018; Ke *et al*. 2020). One *M. citricolor* Illumina short-read assembly is publicly available at NCBI databases (Kotlobay *et al*. 2018). This assembly of *M. citricolor* resulted in a 75.1 Mbp genome comprising over 12,000 contigs (N50: 16,653 bp). Furthermore, 32 genome assemblies for non-pathogenic *Mycena* species are also available in NCBI at different resolution levels, most generated by JGI’s Mycocosm project. Notably, despite these genomic studies within the mycenoid clase, no other studies have delved into exploring the genetic basis for specific pathogenicity mechanisms in *M. citricolor* so far.

In our study, we hypothesised the existence of genome-wide changes, and utilised the lifestyle differences of *M. citricolor* as compared to other saprophytic species in the genus, to identify potential pathogenic genes for further analysis. Our approach involved examining the adaptations that led to the distinct lifestyle of *M. citricolor* and using them to pinpoint a set of candidate genes associated with pathogenicity. For that, we performed a long-read assembly of the highly virulent *M. citricolor* isolate 010 and used in silico techniques to identify virulence-related proteins that are secreted and have extended copy-number variation (CNV) between them. Our aim is to enhance the comprehension of the genetic foundation that facilitates the pathogen’s infection strategy, specifically on coffee plants. This would provide useful insights for conducting further research on the pathogen’s genetic basis, allowing comprehension of its biology, epidemiology, and disease management strategies. Additionally, this comprehensive understanding of the pathogen’s genetic mechanisms can aid in the development of ALS-resistant coffee varieties by incorporating germplasm screening eventually.

## Methods and Materials

### *Mycena citricolor* isolation and cultivation

One isolate of *Mycena citricolor,* named 010, was previously isolated and characterized in FNC-Cenicafé’s Plant Pathology lab (Arciniegas *et al*. unpublished) from leaves affected by American Leaf Spot (ALS) from a *Coffea arabica* cv. Castillo tree in FNC-Cenicafé’s experimental station at El Tambo (Cauca, Colombia). The isolation process for this particular isolate involved several steps. ALS-affected leaves were washed several times with sterile distilled water and placed into humid boxes for 1-2 weeks to stimulate the production of asexual gemmae. Clean healthy gemmae were individually collected and placed on petri dishes with Potato Dextrose Agar (PDA, Sigma Aldrich), and incubated in the dark at 21°C for three weeks. To increase the quantity of required mycelia for DNA extraction, six 0.5 cm agar discs with fungus were transferred into 100 mL of Potato Dextrose Broth (PDB, Sigma Aldrich) and kept in orbital shaking at room temperature for 15 days. After that, the mycelium aggregates were filtered, superficially washed with sterile distilled water, and resuspended into 1X PBS (Phosphate-Buffered Saline). For long-term storage, mycelium was collected in 10 mL tubes, covered with perforated Parafilm® and stored at -80°C for one week, later freeze-dried (lyophilised), weighed inside 1.5 mL cryogenic tubes, and stored at -80 °C until use.

For large number of gemmae production and isolation, the same type of fresh mycelium aggregates produced in PDB mentioned previously were rinsed four times with 100 mL sterile distilled water and placed individually on disinfested detached *C. arabica* cv. Caturra coffee leaves. Leaves with fungus aggregates were incubated in controlled environment cabinets in FNC-Cenicafé Plant Pathology lab at 21°C, 10h low-intensity light photoperiod and ∼99% relative humidity and were frequently sprayed with water. After 1 week, newly developed gemmae were collected when matured (bright yellow colour) by softly shaking each aggregate in 5 mL of ultrapure water. Gemmae were filtered and resuspended in PBS buffer. Gemmae were transferred into cryogenic tubes, freeze-dried, and stored at -80 °C until use.

For basidiocarp large production and isolation, disinfested *C. arabica* cv. Caturra young leaves were inoculated with a suspension of isolated gemmae, as described before. Inoculated leaves were maintained in controlled environment cabinets, as described before. Newly developed gemmae emerged one to two weeks later, and basidiocarps arose one month after inoculation from advanced ALS necrotic lesions. Basidiocarps were collected from the pediceĺs base using a fine clamp, washed, and resuspended in PBS, frozen and freeze-dried, and stored at -80 °C until use.

### Genome sequencing and assembly

DNA and RNA were extracted and sequenced at Novogene (Hong Kong) from shipped freeze dried fungus tissue samples, following regulatory and phytosanitary international procedures. We obtained 2,982,005 reads using four PacBio Sequel Single Molecule Real Time (SMRT) from one DNA sample. We also sequenced RNA by Illumina Platform (PE150), 250∼300 bp insert length, and generated 10.7M, 12.2M and 22.5M reads from each gemmae, mycelia and basidiocarps, respectively. A previous short-reads *Mycena citricolor* genome assembly (Kotlobay *et al*. 2018) was downloaded from NCBI (assembly accession GCA_003987915.1), which corresponded to the *M. citricolor* isolate CBS 193.57, collected in coffee plants in Costa Rica.

We used two long-read assemblers: The diploid-aware assembler FALCON version 20180312 (Chin *et al*. 2016), and CANU version 2.0 (Koren *et al*. 2016). FALCON and CANU were selected because they use different approaches (HGAP vs. MHAP) and are broadly used in fungi assembly. Each assembler was run twice with two different parameter sets. FALCON was firstly run with a minimum read length of 500 bps, “expected genome size” value of 100 Mbps, and the options listed by (Miller *et al*. 2017). This assembly was called “FALCON2”. Another FALCON assembly was generated with the same options but *relaxed overlapping* (pa and ovlp): “-e 0.75” instead of 0.95, “-s 100” instead of 1000, “-h 480” instead of 1024, and “-l 2000” instead of 4000. This assembly was called “FALCON3”. CANU was run with default options and two different expected genome size of 220 Mbps (“CANU1”) or 100 Mbps (“CANU2”).

### Quality check of *de novo* assemblies

The quality of the four long-read assemblies was compared using four tools: BUSCO (Seppey *et al*. 2019) to check for the presence of conserved single-copy orthologs (**Table S2**), QUAST (Gurevich *et al*. 2013) to obtain genome contiguity statistics (**Table S1** and **Figure S1**), KAT (Mapleson *et al*. 2018) to generate Kmer plots to detect any content in the raw reads absent in the assemblies (**Figure 2A**), and D-genies (Cabanettes and Klopp 2018) for whole-genome assemblies to compare whole genome synteny and generate dot plots from whole genome alignments (**Figure 2B** and **2C**). BUSCO version 4.1.2 was used to assess the completeness of the four assemblies of *Mycena citricolor* isolate 010, as well as the *Mycena citricolor* assembly downloaded from NCBI (GCA_003987915), using the databases of single-copy markers from BUSCO, “fungi” and “Agaricales”. Contiguity stats were obtained using QUAST version 4.3 with default parameters, except “min-contig 1000”. Minimap2 was used to generate whole-genome alignments using the option “-DP” to retain secondary alignments. Alignments with scores under 200, a percentage of identity under 50%, and shorter than 5 Kbps were discarded before plotting using D-Genies (http://dgenies.toulouse.inra.fr). Finally, K-mer hashes were generated from each assembly and the corrected reads, and compared using the tool “comp” from the Kmer analysis toolkit (KAT) version 2.3.4 (Mapleson *et al*. 2017). The resulting plot (“Kmer comparison plot”) showed stacked histograms of the Kmer frequency in the reads and divided and coloured by copy number in the assembly (e.g. black for zero copies or absent in the assembly, or red for one copy present in the assembly). In the case of the two assemblies produced with Falcon, we concatenated the primary contigs (p_contigs) and the alternative ones from homologous genomic region (a_contigs).

**Figure 2.**
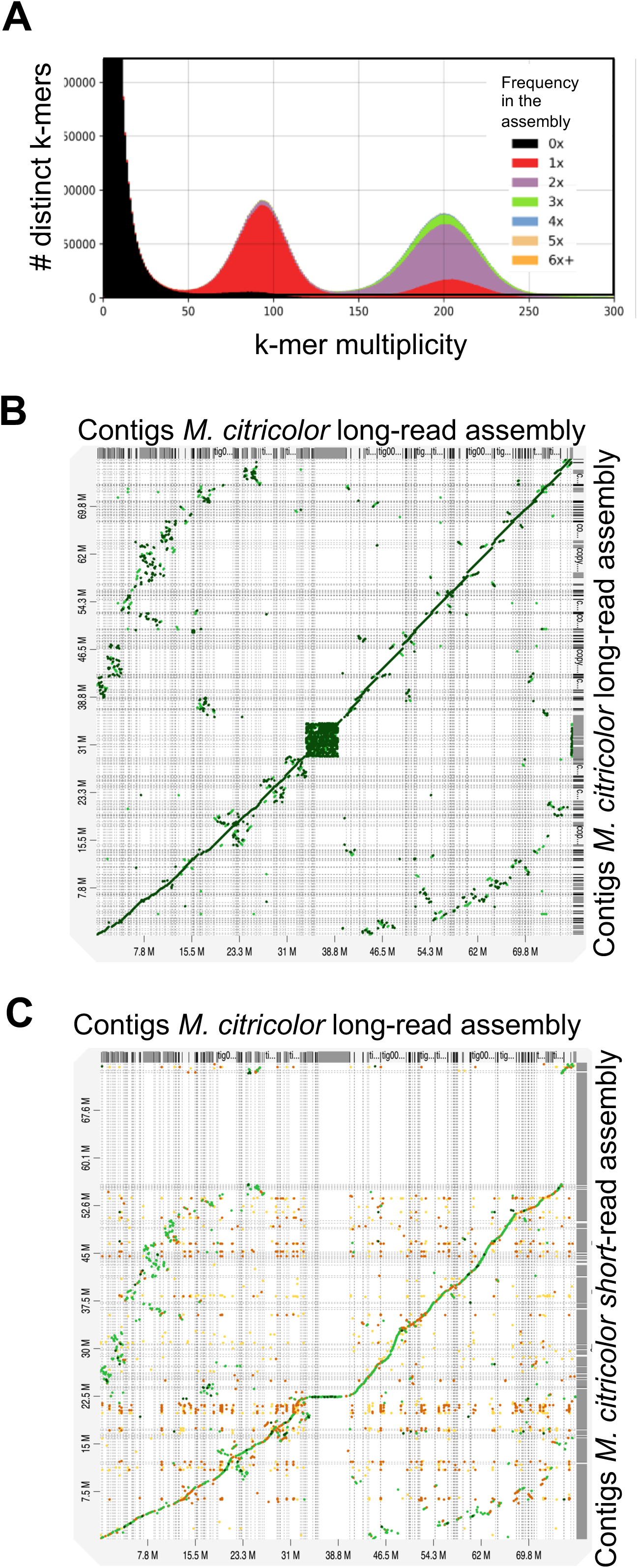
Quality assessment plots for the de-novo hybrid assembly of *Mycena citricolor* (CANU1). A) K-mer spectra comparison plot for *M. citricolor* assembly produced using KAT. Synteny dot plots showing whole genome alignments between *M. citricolor* assembly to itself (B) and to the short-read assembly (C; y-axis *M. citricolor* GCA_003987915.1) produced using D-genies.

### Gene structure and function annotation

Firstly, *ab initio* transcripts were assembled from the RNA-seq generated from three tissues (mycelia, gemmae and basidiocarps) from *M. citricolor* isolate 010. For that, the reads were aligned to the “Myci” genome using HISAT2 v. 2.1.0, with the “dta” option to tag the strand (40). The resulting spliced alignments were used to assemble transcripts with Stringtie v. 2.1.1 (Kovaka *et al*. 2019). We also downloaded all the transcripts and proteins from *Mycena* species available at JGI’s Mycocosms. The Stringtie transcripts (assembled from RNA-seq) and the transcripts and proteins from the other *Mycena* species were used as input to Maker version 2.31.8 (Cantarel *et al*. 2008; Stanke *et al*. 2008), using the default workflow to obtain gene models. Finally, the gene models from MAKER, the Stringtie transcripts, and splice junctions inferred with Portcullis v. 1.0.2 (Mapleson *et al*. 2018) from the HISAT2 spliced alignments were provided to Mikado version 2.0.prc2 (Venturini *et al*. 2018) to generate a final annotation of gene models. The functional annotation based on homology with NCBI’s NR and Interpro and associated GO terms was completed using Blast2GO (Conesa *et al*. 2005).

### Phylogenetic orthology and protein duplication

Phylogenetic analysis based on the prediction of groups of ortholog proteins (orthogroups) was done with Orthofinder version 2.3.12 (Emms and Kelly 2019) and used to locate *Mycena citricolor* in relation to other members of the *Mycena* genus, using the default options of the “MSA” workflow (-M msa). This infers maximum likelihood trees from multiple sequence alignments (MSA) using RAxML with default options for bootstrapping. Proteomes from 26 non-pathogenic *Mycena* species and other species representative of the Agaricales order, were downloaded from NCBI and JGI’s Mycocosm database (Grigoriev *et al*. 2014). Orthofinder also generated gene trees for every orthogroup, a consensus species tree, and identified gene expansion events used in the later candidate gene analysis. Phylogenetic trees were replotted using Dendroscope version 3.7.5 and Figtree v. 1.4.4. Duplicated candidate genes were identified by extracting the genes in orthogroups expanded in the node that divided pathogenic and non-pathogenic *Mycena* species (node 15) in the “species duplications tree” generated by Orthofinder. To test for any bias resulting from the different gene annotation pipelines used (JGI’s annotation pipeline or Maker), Orthofinder was run again (**Figure S3**) after doing new annotations using the same Maker pipeline used for *M. citricolor* (short-read assembly: GCA_003987915) and *Mycena rebaudengoi*. Orthofinder was run a final time including the proteomes of *Botrytis cinerea* and *Sclerotinia sclerotiorum* to identify the orthologues between *M*. *citricolor* and these two species.

### Secretome prediction

The secreted proteins (secretome) from the pathogenic *M. citricolor* strain 010, *M. citricolor* GCA_003987915 and non-pathogenic *M. pura* were determined by identifying proteins with signal peptide sequences but without transmembrane domains. Proteins with signal peptides were determined using TargetP version 2.0 and SignalP version 5.0 (Almagro Armenteros *et al*. 2019; Armenteros *et al*. 2019). Transmembrane helices were identified using TMHMM version 2.0 (Krogh *et al*. 2001; Möller *et al*. 2001). These programmes were run through the online version of The Technical University of Denmark (https://services.heatlhteach.dtu.dk). The lists of proteins from SignalP, TargetP and TMHMM for each analysed *Mycena* proteome were compared to select proteins with a signal peptide but no transmembrane helix domains.

### Homology to experimentally supported virulence factors

To identify virulence factors from the pathogenic *M. citricolor* strain 010, *M. citricolor* GCA_003987915 and non-pathogenic *M. pura*, proteomes were aligned to the Plant Host Interactions database (PHI-base) at http://www.phi-base.org/ (Urban, Cuzick et al. 2020) using DIAMOND (version 0.9.10) (Buchfink *et al*. 2021). PHI-base contains virulence factors that have been experimentally verified, only virulence factors in pathogenic fungi species were used as targets. Alignments were required to have over 50 % identify, a bit score over 50, and be longer than 50 amino acids.

### Expression values (RPKM) from RNA sequencing

RNA-seq data for mycelium, gemmae and basidiocarp from *Mycena citricolor* strain 010 was analysed using Kallisto version 0.46.1 (Bray *et al*. 2016), and the assembled genome and annotation as reference, to estimate RPKM (Reads Per Kilobase Million) and TPM (Transcripts Per Kilobase Million) expression values. TPM values under 1e-6 were rounded to 1e-6. Log2 fold-change was calculated between gemmae and mycelium TPM values, and transcripts with fold-changes greater than 4 were labelled as “over-regulated” or “repressed” in gemmae. Only transcripts with TPM value >= 5 TPM in either gemmae or mycelia were considered.

### Identifying candidate gene clusters in *M. citricolor* strain 010

Gene density was calculated using BEDTools (Quinlan and Hall 2010) in 25 Kbp windows after filtering the genome annotation for secreted, virulent and duplicated proteins. Only genome contigs longer than 500 Kbp were included. Any 25 Kbp windows containing at least three secreted or five virulent proteins were considered as candidate clusters.

## Results

### Sequencing, assembly, and annotation of a reference *M. citricolor* genome

DNA was extracted from a highly virulent *M. citricolor* isolate, named “010”, and sequenced using Pacbio Sequel SMRT sequencing to generate over 2.9 M reads in four flow cells. We later generated four assemblies and compared them: the two CANU assemblies showed longer contigs and greater N50 compared to the two FALCON assemblies (**Table S1**; **Figure S1**). Kmer spectra plots (**Figure 2A**) showed the expected homozygous and heterozygous peaks for a diploid genome and verified Kmer content in the reads absent in the assemblies was very low (black area in **Figure 2A**). This was observed in all the Kmer-based analysis of the four assemblies (Data not shown). However, the CANU assemblies had more “complete” and less “missing” BUSCO single-copy markers than the FALCON assemblies: 92.5-93.7% complete markers compared to 77.3-81.9% using FALCON (**Table S2**). The assembly “CANU1” was eventually selected for downstream analysis and renamed as “Myci”. The final assembly consisted of 493 contigs, N50 was 599,315 bp, with GC content of 53.12%, and the largest contig was 4.15 Mbp. The total length was 77.5 Mbp (**Table S1**).

Whole genome alignments of the new long-read assembly against itself (**Figure 2B**) and the public short-read assembly from the same species, i.e. the two existing *M. citricolor* assemblies (**Figure 2C**) evidenced high repetition in the centromeric region and sporadic conserved homology between some regions (dispersed dots in **Figure 1B**). There were no detectable large chromosomal rearrangements in the genome of *M. citricolor* isolate 010 (**Figure 1B**). Comparison between both isolates evidenced high conservation between them, as well as ∼10Mbp of content in the short-read assembly that was not present in the long-read assembly (**Figure 2C**).

The gene annotation for *M. citricolor* 010 included 21,425 genes and 24,533 mRNAs. Of these, 7,476 could be associated with a function and at least one GO term. Stringtie, using RNA alignments to the reference genome, assembled 27,199, 26,315 and 30,052 transcripts in gemmae, basidiocarps, and mycelial tissue, respectively.

### Phylogenetic orthology of *Mycena citricolor*

A phylogeny (**Figure 3**) was constructed using protein orthogroup inference from the predicted proteomes of 55 Agaricales fungi, representing *M. citricolor,* the saprophytic *Mycena* spp. and the wider Agaricales order, using the OrthoFinder tool (Emms and Kelly 2019). *M. citricolor* isolate 010 clustered closely to *M. citricolor* GCA_003987915.1 in a different branch than the rest of the non-pathogenic *Mycena* spp.

**Figure 3.**
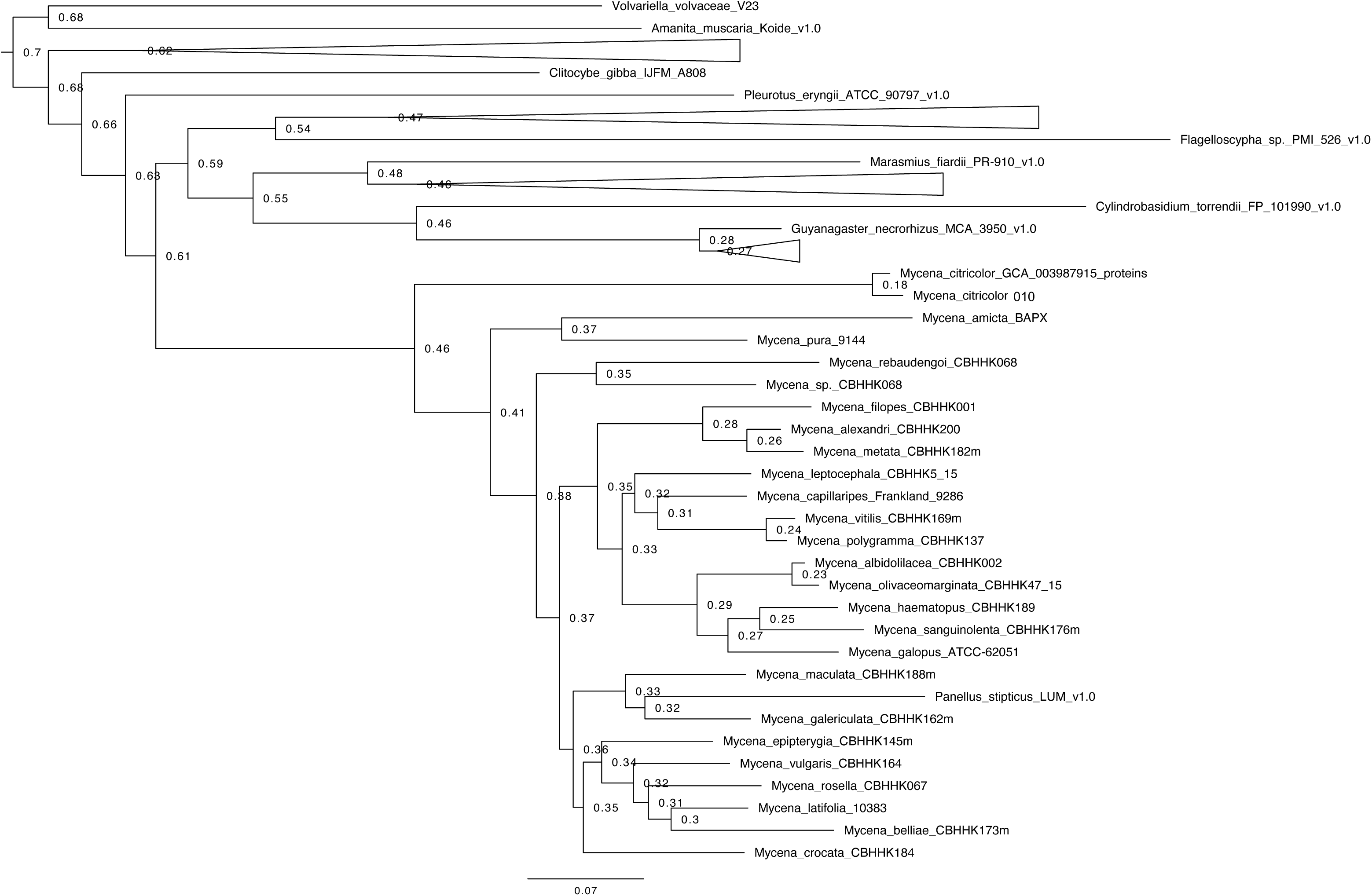
Phylogeny derived from orthologous genes in the predicted proteomes of 55 representative fungi members in the Agaricales order, including *Mycena* genus members, along with *M. citricolor* 010.

Orthofinder also identified gene duplication events in each node (**Figure S2**). Focusing on the node in the tree where pathogenic and non-pathogenic *Mycena* spp. diverged, we identified 620 orthogroups whose protein members were classified as “gene duplication events” by orthofinder, which included 1,987 proteins from our *M. citricolor* isolate 010 long-read genome assembly and 2,067 proteins from the public *M. citricolor* short-read assembly (**Table S3**).

Finally, to verify that the split between pathogenic and non-pathogenic was not a consequence of differences in annotation pipelines, a new run of Orthofinder was done where we included new annotation of both *M. rebaudengoi* and *M. citricolor* GCA_003987915 using our gene annotation pipeline (**Figure S3**), which verified the previous results with 55 individuals.

### Secretome prediction and homology to experimentally supported virulence factors

The proteomes of *M. citricolor* 010 (21,425 proteins; long-read assembly), *Mycena citricolor* GCA_003987915 (26,793 proteins; public short-read assembly), and the non-pathogenic *M. pura* isolate 9144 (28,619 proteins; from JGI’s Mycocosm) were used to infer proteins with signal peptides but not to be trans-membrane helices (non-TMH). *M. pura* was randomly selected among the non-pathogenic *Mycena* species. 1,740 secreted genes were identified for *M. citricolor* 010 (∼8.1%), 1,995 secreted genes in *M. citricolor* GCA_003987915 (∼7.4%) and 2,131 proteins (∼7.4%) in *M. pura*. E.g., these 1,740 correspond to non-TMH proteins identified by either TargetP (357) or SignalP (5), or both (1378) in *M. citricolor* 010 (**Figure S4**; **Table S4**). Proteins with signal peptides were determined using TargetP version 2.0 and SignalP version 5.0 (Almagro Armenteros *et al*. 2019; Armenteros *et al*. 2019). Transmembrane helices were identified using TMHMM version 2.0 (Krogh *et al*. 2001; Möller *et al*. 2001).

All proteins were then compared to the plant host interaction database (PHI-base), which contains experimentally verified virulence factors (Cuzick et al. 2020). The analysis showed that 746 genes were homologous to virulence factors from *M. citricolor* 010 isolate, 957 genes were homologous to virulence factors from *M. citricolor* GCA_003987915 (∼3.6%), and 556 genes homologous to virulence factors from non-pathogenic *M. pura* (∼1.9%), all aligned with experimentally verified fungal virulence factors in the database (**Figure S4**; **Table S5**). Copy-number variation based on homology to similar virulence factors was calculated for *M. citricolor* and *M. pura* as well (**Table S6**).

Enrichment analysis of Gene Ontology (GO) Terms among either the secreted proteins or the experimentally-verified virulence factors showed minor differences between the two pathogenic *M. citricolor* and the saprophytic *M. pura* isolates: GO terms with much lower p-values in both pathogenic isolates, but not the saprophytic were: sucrose and starch metabolism, protein dephosphorylation, glycosylation, pantothenate and (iso)leucine biosynthesis, and cellular oxidant detoxification (**Figure S5**).

### Tissue-specific TPM comparison

TPM (Transcripts per Kilobase Million) expression values were obtained from gemmae and mycelia tissue from isolate 010. Transcripts were labelled as “over-expressed” or “repressed” in gemmae if the log2 fold-change was four times in gemmae during plant interaction than in mycelia during *in vitro* liquid culture media growth, resulting in 1,354 (6.3%) and 1,058 (4.9%) transcripts identified in each group, respectively (**Table S7**).

### Co-location of candidate genes

Using the physical locations in the genome of *M. citricolor* 010 of the secreted and virulence factor proteins, we also identified 21 clusters containing at least three virulence factors or five secreted proteins based on PHIbase and Secretome databases (**Table S8**).

## Discussion

We completed the assembly and functional annotation of a high-quality reference genome (**Figure 2**) from an experimentally verified highly virulent isolate of *Mycena citricolor,* named 010 (Angel and Arciniegas, 2023), belonging to the only reported pathogenic species in its genus*. M. citricolor* is the causative agent of ALS in plant hosts, such as coffee (**Figure 1**). A previous unanchored short-read assembly from a different isolate of the same species, collected from coffee plants in Costa Rica, was also used (GCA_003987915.1). Additionally, JGI published several genomes from multiple saprophytic species in the *Mycena* genus. The *M. citricolor* genome was assembled into ∼77.5 Mbp (**Table S1**), which is at the low end of the genus: *Mycena* species genome varied from 62.8 Mbp in *M. haematopus* to 501 Mbp in *M. galopus,* the largest genome reported for the Agaricales fungi order.

Phylogenetic analysis based on orthogroups created from whole-genome proteomes showed that *M. citricolor* branched off from all saprophytic *Mycena* spp. (**Figure 3**), indicating that the divergence in lifestyle is reflected in large molecular functional changes throughout the genome. These result from differences in the acquisition of genetic traits associated with the new pathogenic lifestyle, e.g. the activation of virulence factors, suppression of host defences, alteration of metabolisms, and enhanced nutrient acquisition (Delaye *et al*. 2013; De Silva 2017). We leveraged these genetic changes as a means to identify candidate genes.

Firstly, we identified proteins in orthologous groups with expanded copy-number in *M. citricolor* (**Table S3**). This was possible because *M. citricolor* branched off alone from all saprophytic *Mycena* spp. (**Figure 3**), so we searched for gene copy-number variation (CNV) between branches in the split phylogenetic node. CNV is a common strategy for new function acquisition, and the expansion of pathogenic proteins has been reported in *Colletotrichum* spp., another large genus with numerous species and complex taxonomy, with plant pathogenic, non-pathogenic and saprophytic species (Gan 2016).

Secondly, we identified secreted proteins (**Table S4**) using a genome-wide computational prediction approach (Krogh *et al*. 2001; Möller *et al*. 2001; Almagro Armenteros *et al*. 2019; Armenteros *et al*. 2019). We did not find an increase in secreted proteins in pathogenic *M. citricolor* compared to the saprophytic control *M. pura* (7-8% in both lifestyles), nor a change in the functions of these secreted proteins, as evidenced by similar GO terms. The proportion of secreted proteins did not show a good discrimination between pathogenic and non-pathogenic. However, being secreted and duplicated (increased CNV) was different between pathogenic (∼15% of the secreted were increased in copy-number), compared to non-pathogenic *M. pura* (∼6%) (**Figure 4A-C**). This corresponded to 220 genes (218+2) in the long-read assembly and 231 genes (228+3) in the short-read assembly (**Figure 4A-B**) that were shortlisted as candidates.

**Figure 4.**
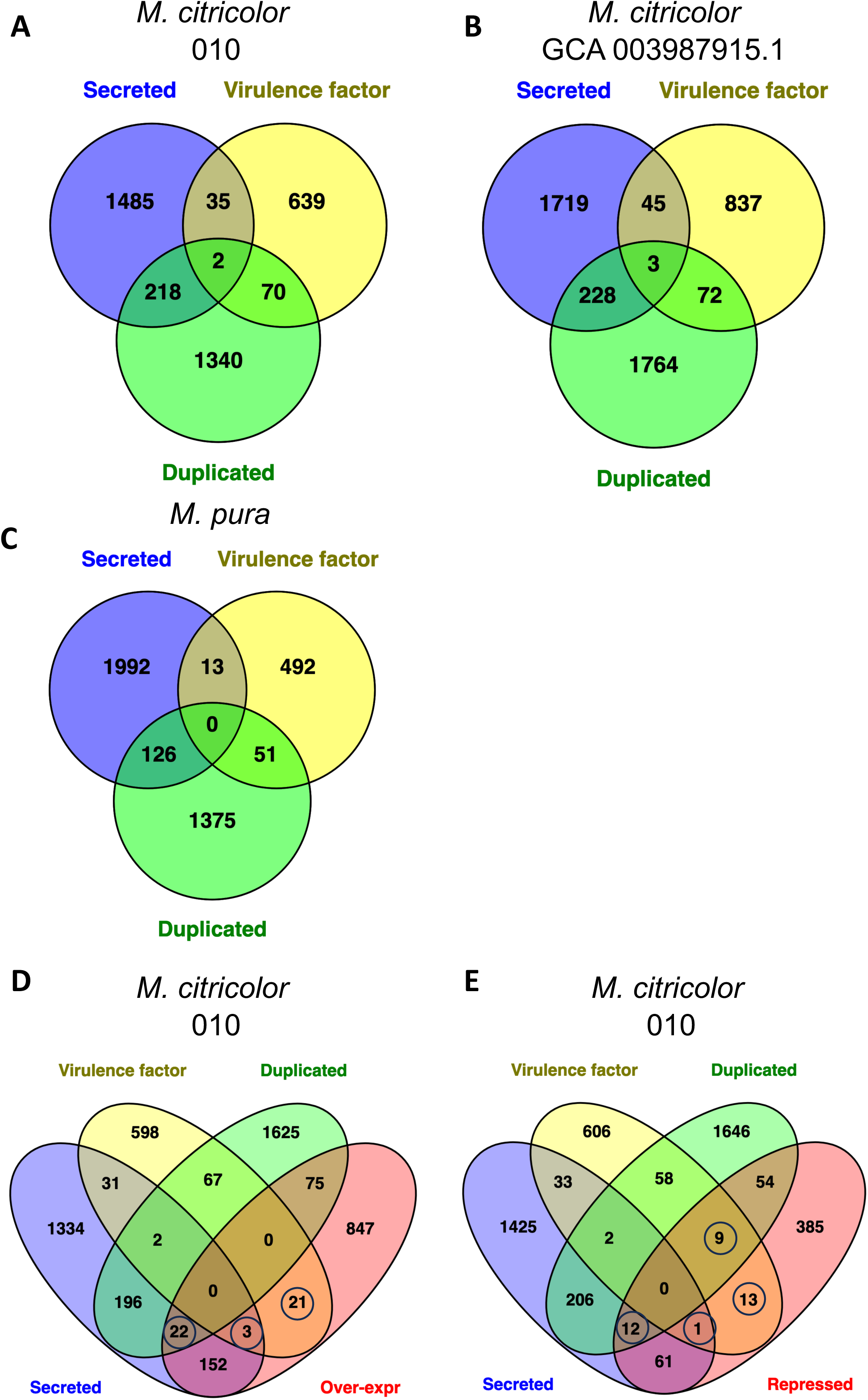
Ven diagram illustrating the number of secreted proteins, virulence factors and duplicated proteins from *M. citricolor* 010 (A), *M. citricolor* GCA_003987915 (B) and the non-pathogenic *M. pura* (C). Additionally, the comparison of all *M. citricolor* 010 proteins to ‘over-expressed’ (D) and ‘repressed’ (E) transcripts is presented (refer to **Table S7**).

Third, we used homology-based annotations, which showed a good discriminator between pathogenic and non-pathogenic species. We found an increase of homologous to verified virulence factors *in M. citricolor* (3.4%-3.6%) compared to *M. pura* (1.9%). We also used number of copies aligning to each target virulence factor to identify any increase in copy-number, resulting in 72 and 75 candidates in each *M. citricolor* assembly that were shortlisted as candidates. The target species were also different, over 5% of the *M. citricolor* queries aligned to a *Botrytis cinerea* homolog as the top result, but not a single protein from *M. pura* aligned to a *B. cinerea* target (**Table S5**). Because of this observation, we annotated every ortholog between *M. citricolor* and *B. cinerea* (**Table S9**). *M. citricolor* obtains nutrients by penetrating the plant tissue as a hemibiotroph, and later kills the host cells by secreting lytic enzymes and organic compounds, such as oxalic acid (Rao and Tewari 1987), causing cell death and tissue decay, as a necrotrophic pathogen. This strategy, switching from a biotrophic to necrotrophic lifestyle, and penetration mode is similarly found in *B. cinerea*, *Tilletia indica, Clarireedia jacksonii*, *Sclerotinia homeocarpa*, and *Sclerotinia sclerotiorum*, which cause economically important diseases in crops worldwide (Giraldo and Valent 2013; Kabbage *et al*. 2015). *S. sclerotinium*, and *Tilletia indica* also secrete oxalic acid to suppress the oxidative burst of the host plant (Cessna *et al*. 2000a; Venu *et al*. 2009; Pandey *et al*. 2018; Townsend *et al*. 2020).

Finally, we used gene expression between *M. citricolor* gemmae and mycelium tissues to identify genes regulated during infection in the fungus on inoculated coffee leaves, but not in mycelia collected from *in vitro* cultures in liquid media. Because we had no replicates, we opted for a fundamental analysis where we quantified normalised expression as TPM per sample, and identified genes strongly regulated during infection. We defined strong regulation as a log2 fold-change over 4 (either positive or negative) between gemmae during plant interaction and mycelia growing in liquid media. Approximately 11% of the genes showed strong regulation under these criteria.

### Selection of candidate genes

In summary, two criteria discriminated between pathogenic and non-pathogenic: proportion of homologous genes to virulence factors, and proportion of genes secreted and expanded in number of copies (copy-number). To further narrow down the candidate genes, we additionally focused on genes strongly regulated in gemmae during plant interaction in inoculated coffee leaves (**Figure 4**). This limited our candidates to genes from our *Mycena citricolor* isolate 010 genome assembly, where gene expression data was available. **Table S9** incorporated all previous gene annotation information for the 21,425 genes in this assembly.

35 virulence factors were strongly regulated (FC >4) in gemmae during infection (**Table 1**). Table 1 summarises their presence in related plant pathogenic fungi and pathogenic effect based on the information in the PHI-base database. All these listed genes could be considered essential for pathogenicity in *M. citricolor*. We observed a dramatic expansion in copy-number in the well-known virulence factors CUTA (x4.6), MET3 (x4), XYN11 (x4), NoxB (x3.3) (Chakdar *et al*. 2019), MFS1 (Pereira *et al*. 2013), CUTA (Gupta *et al*. 1995) and NoxA (Zhu *et al*. 2021) and NoxB. As well as expanded, these were highly regulated in gemmae and showed high identity to proteins from *B*. *cinerea, F. oxysporum* or *S. sclerotiorum* pathogenic fungi with some similarities in lifestyles (Williamson *et al*. 2007; Segmuller *et al*. 2008; Morita *et al*. 2013; Nordzieke *et al*. 2019). It has been recently highlighted in the literature that the early stages of adhesion and penetration in these pathogens signify a transient biotrophic phase, succeeded by the deployment of necrotrophic strategies aided by oxalic acid and various virulence factors (Kabbage *et al*. 2015; Liang and Rollins 2018; Newman *et al*. 2023). In biotrophic and hemibiotropic pathogenic fungi, the secreted effectors could trigger resistance or immune responses (Giraldo and Valent 2013). In contrast, in necrotrophic fungi, secreted effectors contribute to susceptibility, causing tissue weakening and cell death to continue their life cycle (Liang and Rollins 2018).

**Table 1:**
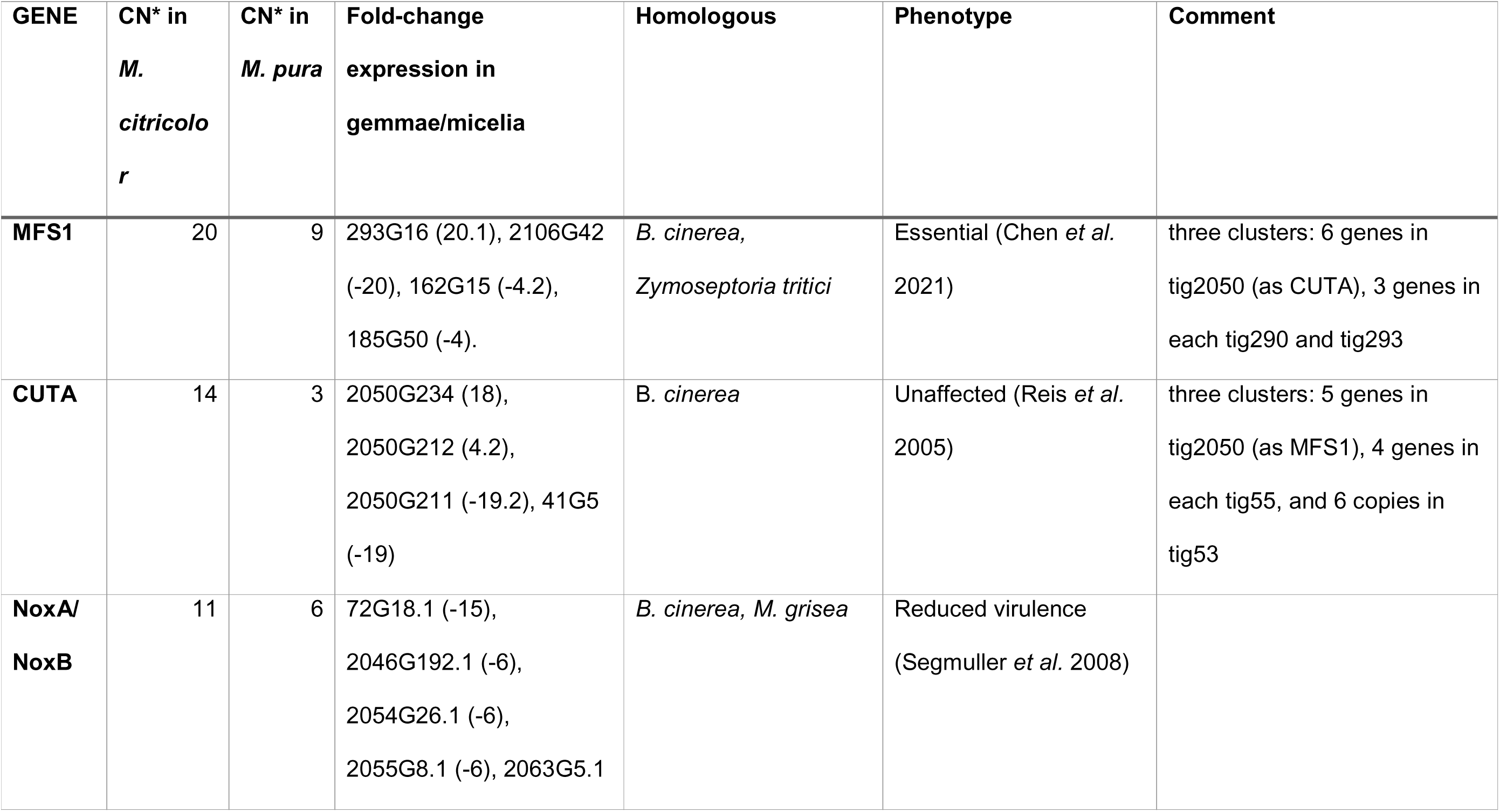
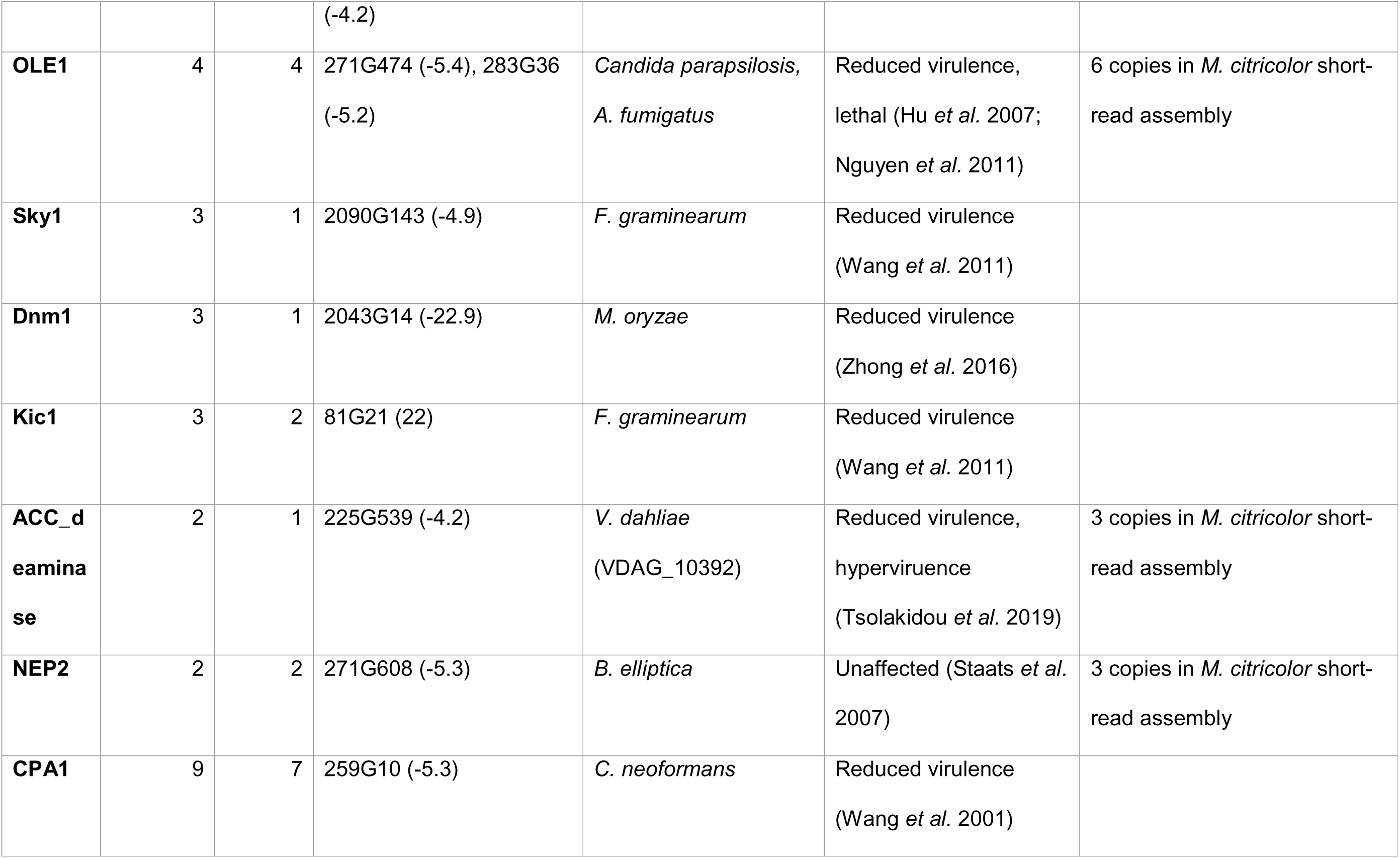
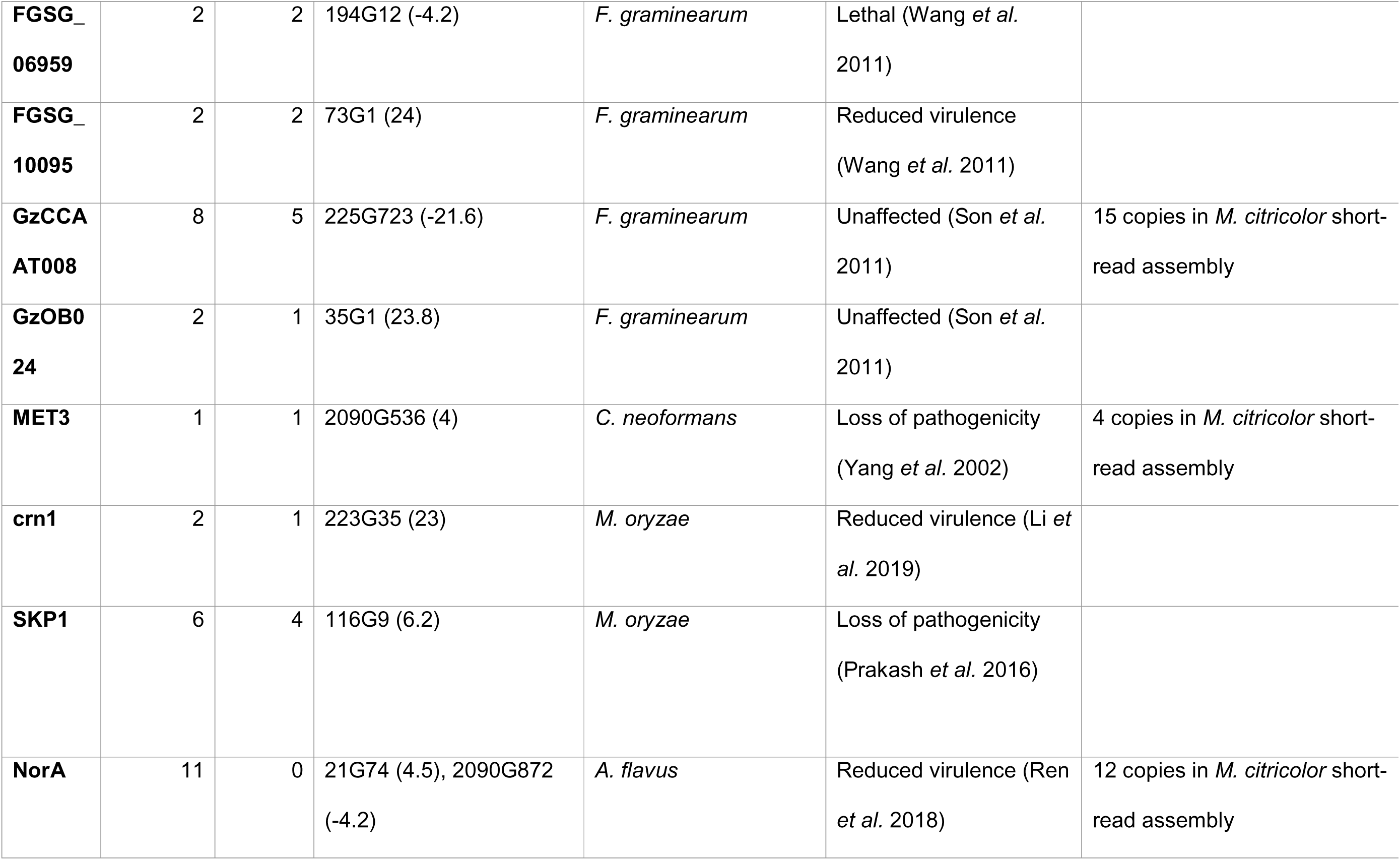
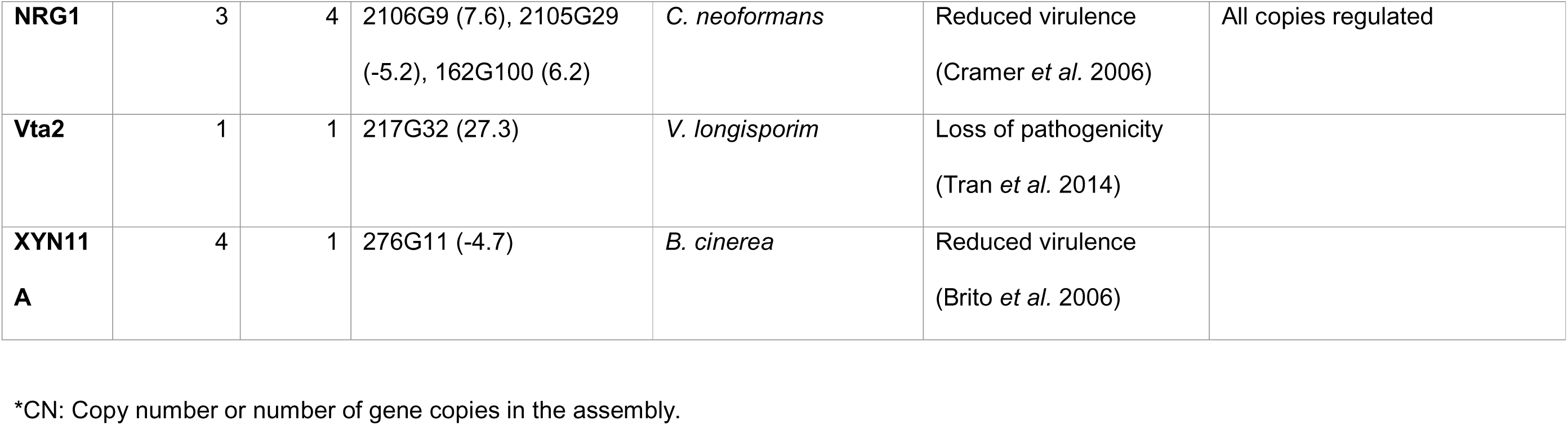
Summary of 35 virulence factors identified in *M. citricolor* 010 related to plant pathogenic fungi.

We observed 220 additional genes that were both secreted and expanded in copy number (**Table S9**), but their function was unknown or poorly annotated so far. Among these, 22 genes were strongly regulated and were listed in **Table 2**: seven were homologous to virulence factors if we relax the filtering criteria, including CYP1/5 (two copies), Cda1/5 (two copies) and PsGH7, KRE6 and SAP5 (one copy each); another seven were homologous to each other and members of orthogroup OG350. This orthogroup is well conserved only in the Mycenaceae family. The protein has unknown function. Several of the remaining candidates were also conserved only at the family or genus level (**Table 2**). We identified two regulated Zn(2)-Cys(6) transcription factors exclusive to *Mycena* spp; 2048G136 (-22.6 fold-change) and 271G454 (-4.9 fold-change). Further studies will be required to determine specific roles and processes involving these candidate genes.

**Table 2:** Twenty-two candidate genes in *M. citricolor* 010.

| GENE | ORTHOGR<br>UP | Expression in<br>gemmae | Function | Homologous | Phenotype |
| --- | --- | --- | --- | --- | --- |
| 136G5 | OG552 | 6.1 | Unknown | Ascomycetes |  |
| 15G13 | OG575 | 22.6 | Unknown | *Mycena* spp. |  |
| 185G13<br>5 | OG350 | 26.5 | Unknown | Mycenaceae<br>family |  |
| 19G13 | OG350 | -7.2 | Unknown | Mycenaceae<br>family |  |
| 19G15 | OG350 | -4.3 | Unknown | Mycenaceae<br>family |  |
| 2046G5<br>0 | OG350 | -6.1 | Unknown | Mycenaceae<br>family |  |
| 2046G5<br>1 | OG350 | -7.4 | Unknown | Mycenaceae<br>family |  |
| 2046G5<br>3 | OG350 | -5.9 | Unknown | Mycenaceae<br>family |  |
| 2048G1<br>36 | OG68 | 5 | CYP5/CYP1 | *V. dahliae* | Loss of pathogenicity (Zhang *et al.* 2016) |
| 2067G9<br>5 | OG350 | 29.7 | Unknown | *Mycena* spp |  |
| 2109G1<br>3 | OG176 | -22.6 | Zn(2)-C(6) transcription<br>factor | *Mycena* spp |  |
| 2112G2<br>67 | OG68 | -4.9 | CYP5/CYP1 | *V. dahliae* | Loss of pathogenicity (Zhang *et al.* 2016) |
| 2112G3<br>48 | OG565 | 5.5 | KRE6 | *C. graminicola* | Reduced virulence (Oliveira-Garcia and<br>Deising 2016) |
| 2124G1<br>65 | OG439 | 19.6 | PsGH7a | *P. sojae* | Reduced virulence (Tan *et al.* 2020) |
| 2126G2<br>30 | OG552 | 6.6 | Unknown | Ascomycetes |  |
| 225G56<br>7 | OG311 | 18.6 | Cda1/Cda5 | *U. maydis* | Reduced virulence (Upadhyia *et al.* 2018) |
| 239G22 | OG311 | 4.4 | Cda1/Cda5 | *U. maydis* | Reduced virulence (Upadhyia *et al.* 2018) |
| 271G29<br>4 | OG500 | 4.2 | exo-glucanase | Ascomycetes |  |
| 271G45<br>4 | OG271 | 4.8 | Zn(2)-C(6) transcription<br>factor | *Mycena* spp |  |
| 301G19 | OG879 | 16.7 | SAP5 | *Candida*<br>*albicans* | Reduced virulence (Sanglard *et al.* 1997) |
| 45G27 | OG2608 | 21.2 | Peptidase | Mycenaceae<br>family |  |
| 67G24 | OG2576 | 4.2 | Carboxypeptidase | Ascomycetes |  |

## Conclusions

Our study highlights how a pathogenic lifestyle has shaped the genome of *M. citricolor*. We exploited these genome-wide effects to shortlist over 300 candidate genes using criteria we found to be different between *Mycena* lifestyles. After analysing the genes among these candidates that are regulated during plant interaction in inoculated coffee leaves, we found that most of them were gene copies of 42 known virulence factors. Most were in multiple copies, and some were expanded up to four times in the pathogenic species. This suggests that *M. citricolor* infection relies on molecular processes that are similar to other fungi with comparable pathogenic strategies and lifestyles. This study aims to utilise existing knowledge to expand the genetic characterisation of multiple *M. citricolor* isolates from coffee, as well as a diverse range of cultivated and non-cultivated hosts from different environments and geographical origins. Future studies can help us better understand the differences in biology, epidemiology, ALS disease dynamics and impacts, and effects of climate change. These studies will also aid in developing strategies for integrated disease management and breeding of coffee ALS-resistant cultivars.

## Conflict of Interest

The authors declare no conflict of interest.

## Author Contribution

NCAB and CAAC contributed to the project conceptualisation, funding acquisition, data analysis, and fungal and DNA/RNA sample isolation, preparation, and submission. FDP contributed to the funding acquisition and experimental design. NLLM and JJDV performed computational analysis, experimental design, and funding acquisition. All authors contributed and approved the manuscript.

## Funding

Funding for sampling, sequencing and characterisation was provided by FNC – Cenicafe (Manizales, Colombia), project PAT102001 to NCAB and CAAC, followed by agreement CN-2019-1532. NLLM, JJDV and FDP were supported by the Biotechnology and Biology Sciences Research Council (BBSRC)’s Global Challenge Research Fund BB/P028098/1 (Grow Colombia) and from the BBSRC Core Strategic Programme Grant (Genomes to Food Security) BB/CSP1720/1 and its constituent work package BBS/E/T/000PR9818 (Signatures of Domestication and Adaptation).

## Acknowledgements

We express our gratitude to FNC and Cenicafe for scientific, logistic, funding, and administrative support. We are also thankful to the multiple semestral intern students from the learning agreement between SENA and FNC – Cenicafe, for their logistic support during the experiments. We are also grateful to MSc. Carlos E. Maldonado (FNC – Cenicafe-Plant Breeding Dept.) for his technical suggestions and assistance with sequenced data handling, and Cenicafe’s Plant Pathology Dept.’s faculty and staff for their valuable assistance. Lastly, we want to extend our gratitude to Dr. Alvaro L. Gaitan, FNC - Cenicafe Director, for his leadership.

## Data Availability Statement

Data used in this manuscript have been deposited as study PRJEB69683 in the European Nucleotide Archive.

## Description of the Supplementary Figure and Tables

**Figure S1.** Comparison of sorted contig lengths among four de-novo assemblies (CANU1, CANU2, FALCON2 and FALCON3) using QUAST.

**Figure S2.** Identification of gene duplication events at nodes within the *M. citricolor* 010 phylogenetic orthology (refer to Figure 3) using orthoFinder.

**Figure S3.** Phylogeny derived from orthologous genes in the predicted proteomes of 55 representative fungi members in the Agaricales order, including members of *Mycena* genus, with the de-novo assembled genome and predicted proteome for *M. rebaudengoi*.

**Figure S4.** Ven diagrams illustrating the number of secreted proteins identified via TargetP, SignalP, or TMHMM in *M. citricolor* 010, *M. citricolor* GCA_003987915 and the non-pathogenic *M. pura*, comparing with the PHI-base plant host interaction database.

**Figure S5.** Enrichment analysis of Gene Ontology (GO) Terms among secreter proteins (A) and experimentally verified virulence factors (B) between *M. citricolor* 010, *M. citricolor* GCA_003987915 and the non-pathogenic *M. pura*.

**Table S1.** Quality assessment stats for the four de-novo assemblies, produced with either canu (CANU1 and CANU2) or falcon (FALCON3 and FALCON4) hybrid assemblers, using QUAST.

**Table S2.** Quality and completeness assessment values for the four de-novo assemblies for *M. citricolor* 010 and short-read genome of *M. citricolor* GCA_003987915.1 using the Benchmarking Universal Single-Copy Orthologs tool (BUSCO).

**Table S3.** List of gene duplication events at each node of the *M. citricolor* 010 phylogenetic orthology (refer to **Figure S2**) using OrthoFinder.

**Table S4.** List of secreted proteins identified using tools TargetP, SignalP, or TMHMM in *M. citricolor* 010, *M. citricolor* GCA_003987915 and the non-pathogenic *M. pura*, and comparing them to the PHI-base plant host interaction database (refer to **Figure S4**).

**Table S5.** List of secreted proteins that aligned with verified fungal virulence factors in the PHI-base (refer to **Figure S4**).

**Table S6.** Gene copy-number variation based on homology to similar virulence factors was calculated for *M. citricolor, M. citricolor* GCA_003987915 and *M. pura* (refer to **Table S5**).

**Table S7.** Transcripts per Kilobase Million (TPM) expression values from ‘over-expressed’ or ‘repressed’ genes in gemmae and mycelia from *M. citricolor* 010.

**Table S8.** List of genes contained in 21 clusters containing at least three virulence factors in the genome of *M. citricolor* 010.

**Table S9.** Summary of gene annotation features from 21,425 genes in *M. citricolor* 010.

## Notes

https://www.ebi.ac.uk/ena/browser/view/PRJEB69683

### Competing Interest Statement

The authors have declared no competing interest.

