## Supplementary material for "Genome-wide characterisation of pathogenicity-related proteins in *Mycena citricolor,* the causal agent of the American Leaf Spot in coffee": Suppl. figure S1

Fig S1: Comparison contigs length, sorted from longest from shortest, in four assemblies of the same reads.

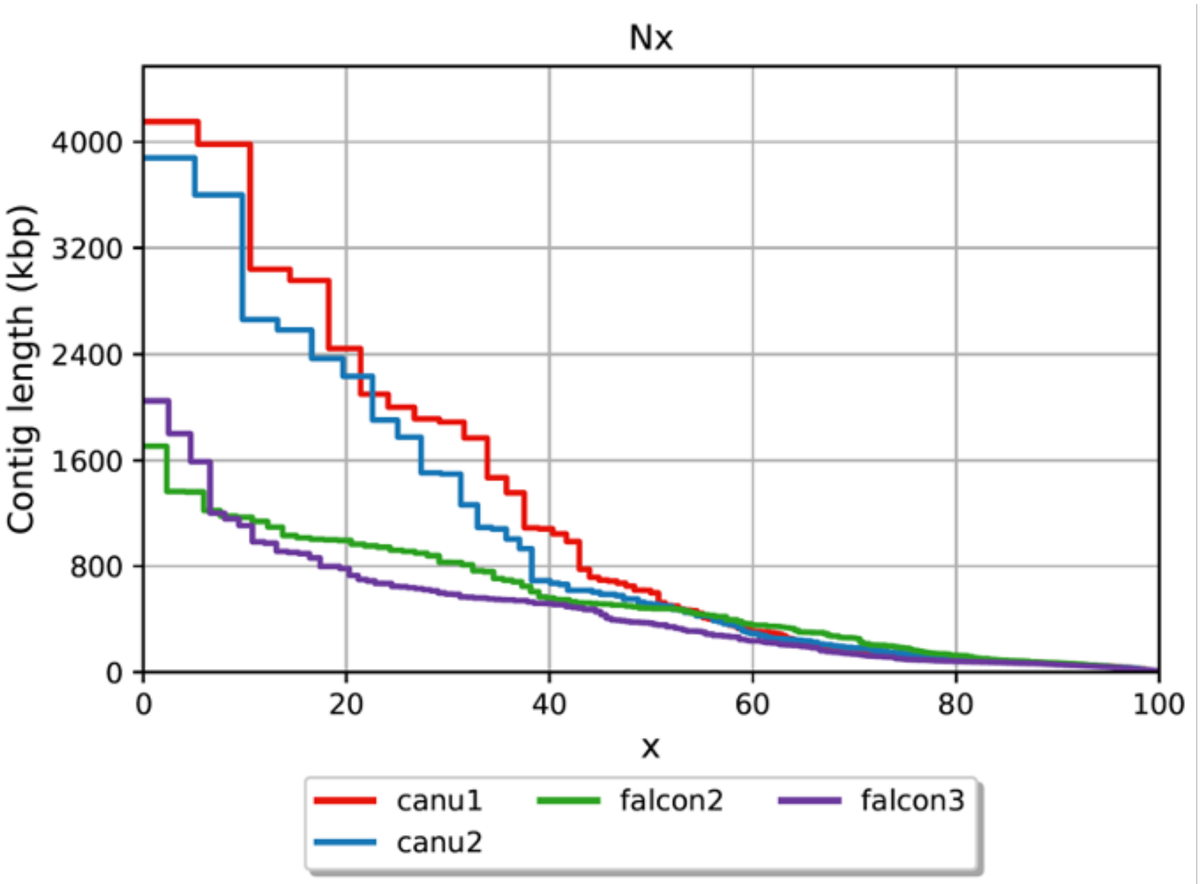
