## Supplementary figures and images for "Genome-wide characterisation of pathogenicity-related proteins in *Mycena citricolor,* the causal agent of the American Leaf Spot in coffee"

### Suppl figure S4

#### *M. citricolor "010"*

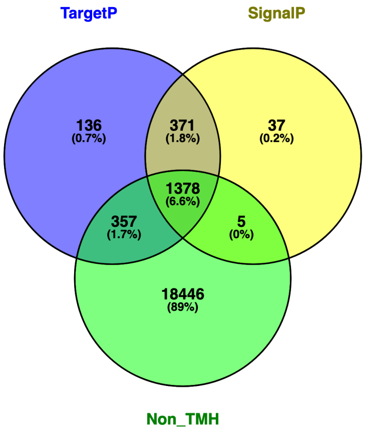


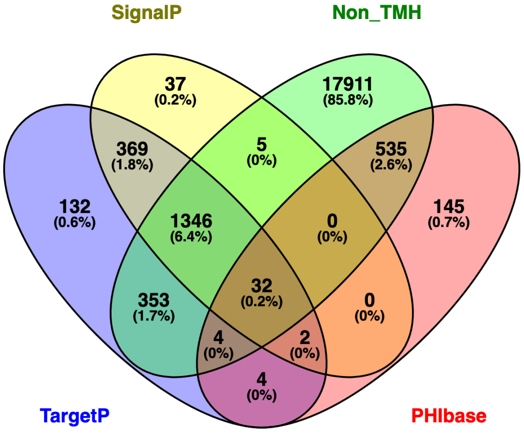


#### *M. citricolor "GCA_003987915"*

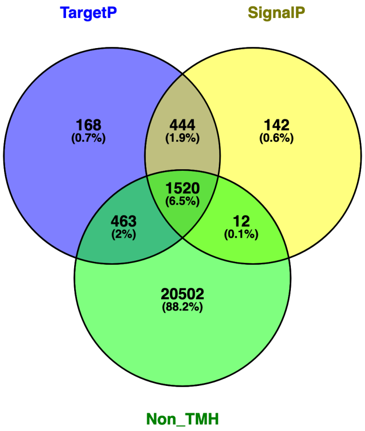


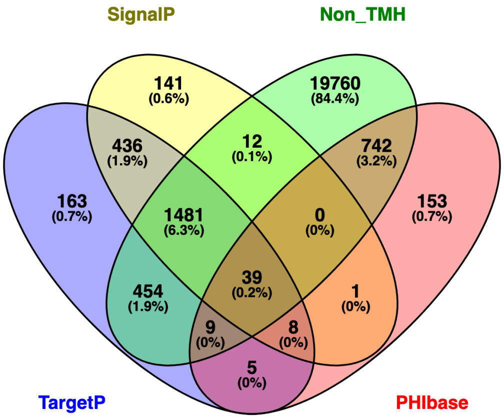


#### *M. pura (non-pathogenic)*

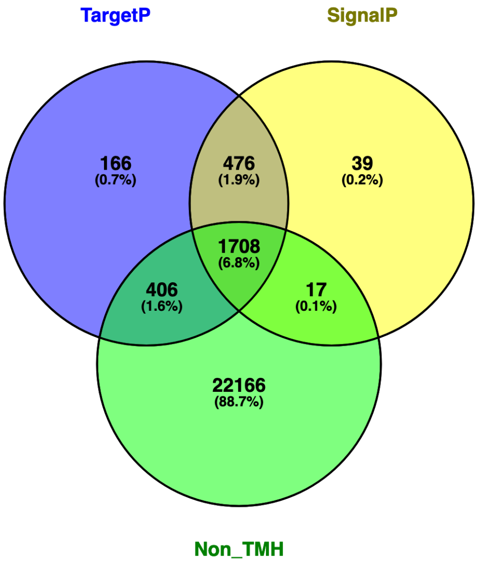


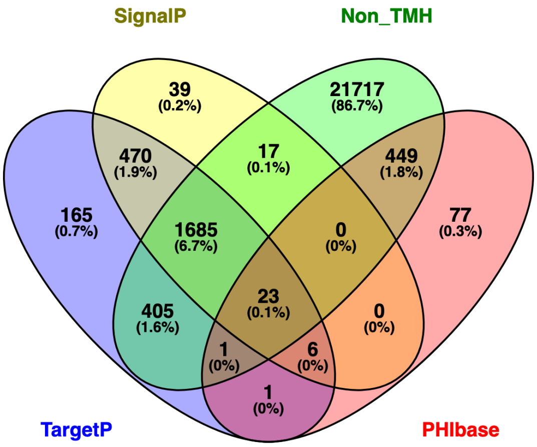

### Suppl. figure S2

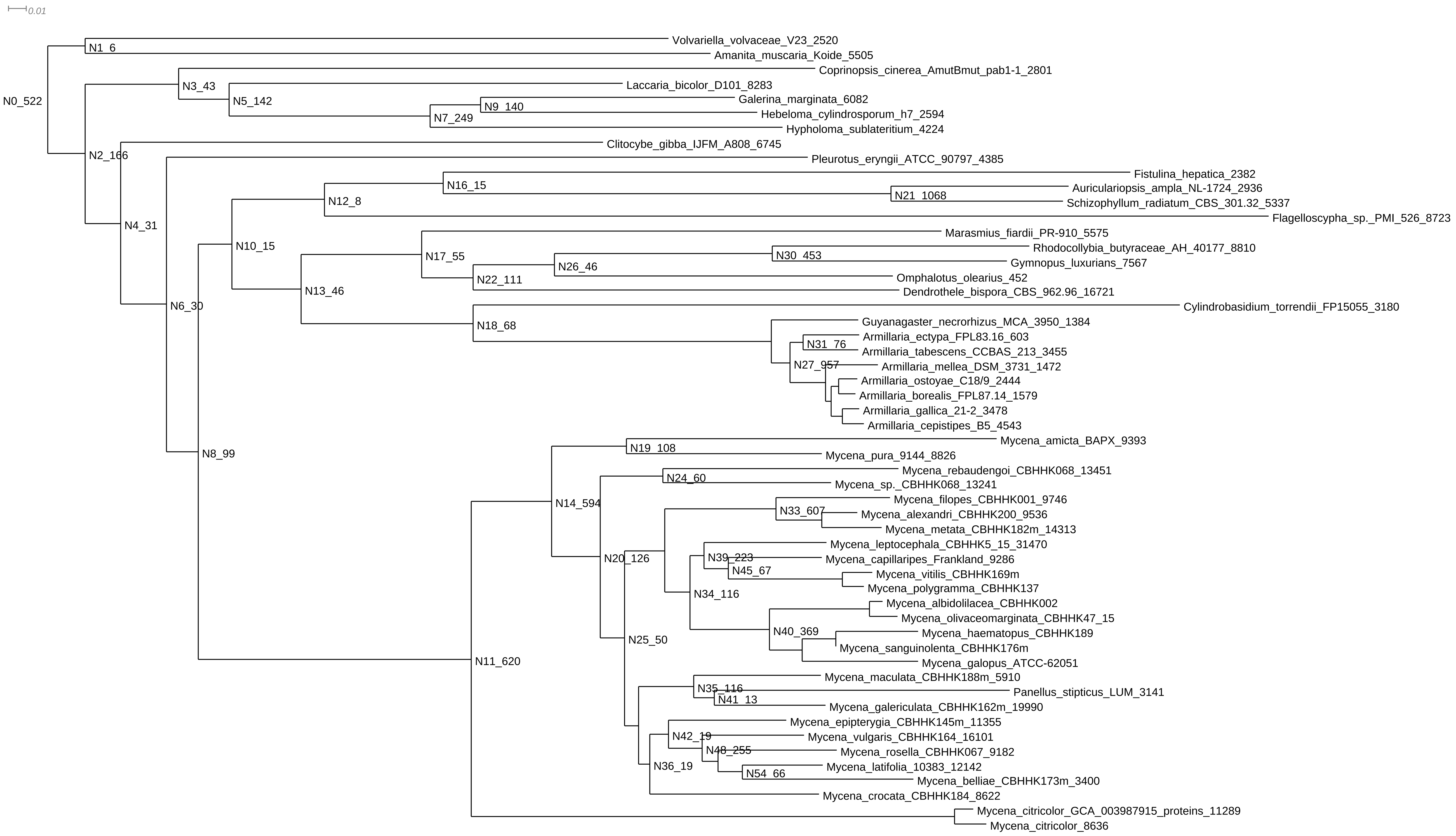

### Suppl. figure S3

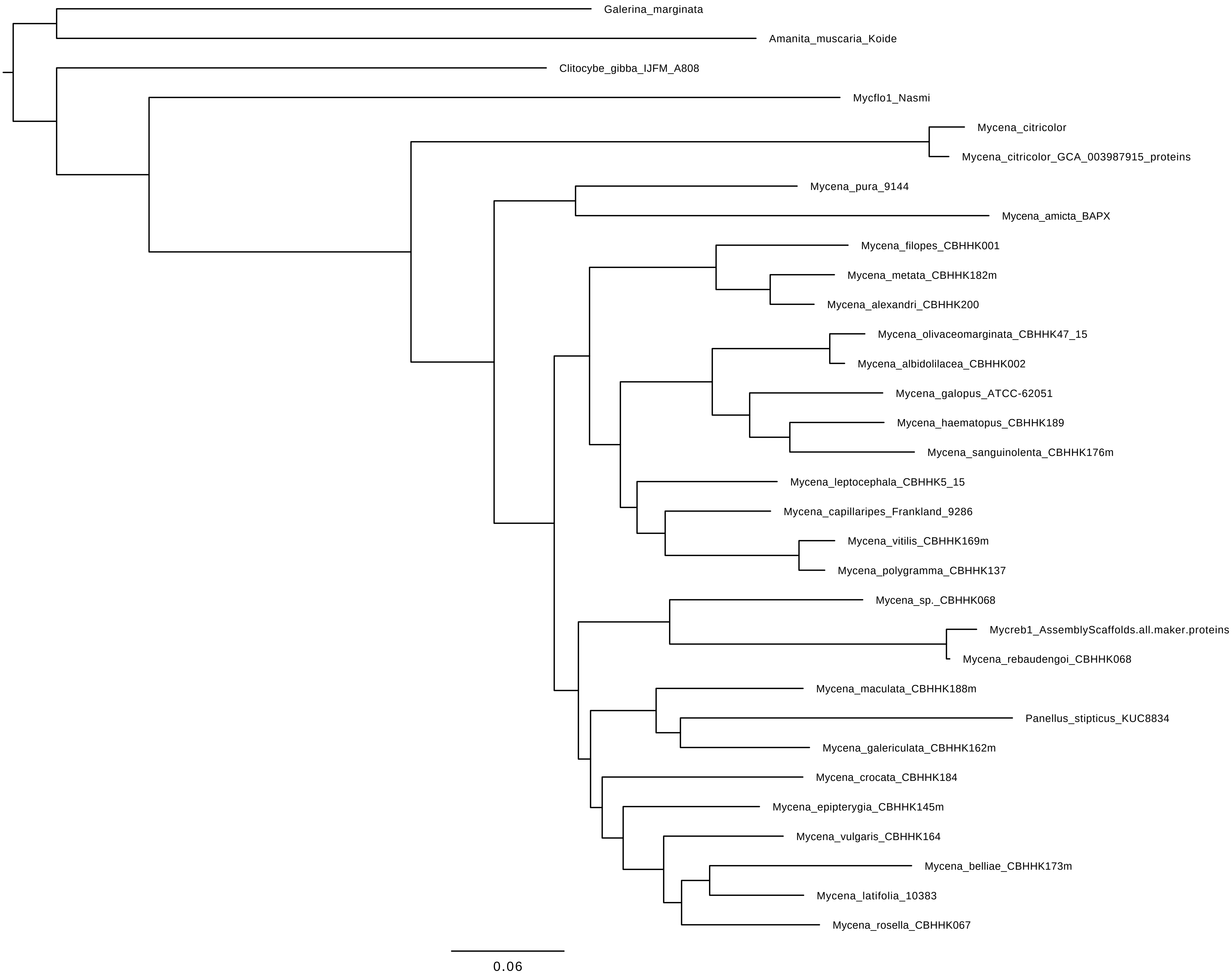

### Suppl. figure S5

##### A: Secreted proteins

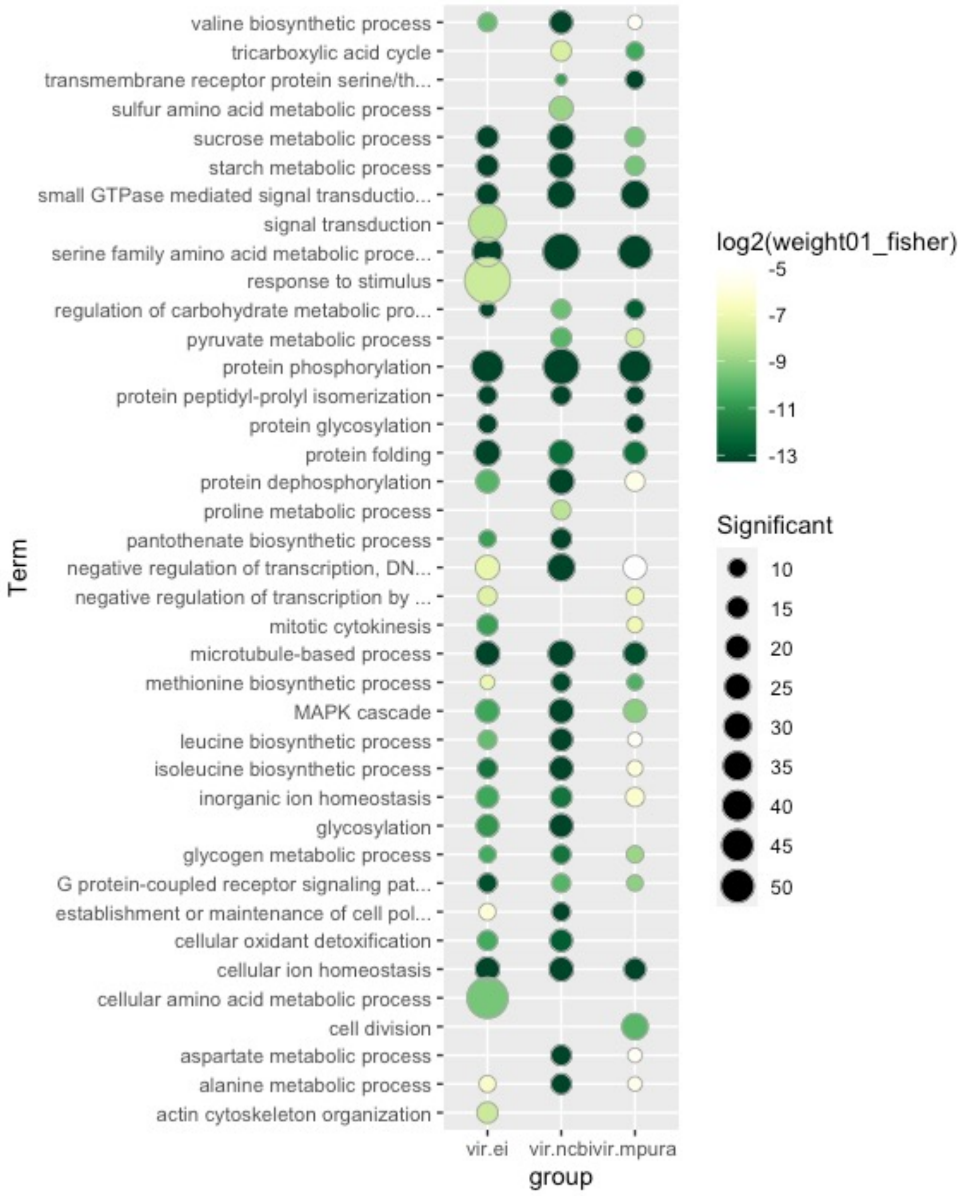


##### B: Virulence-factors secreted proteins

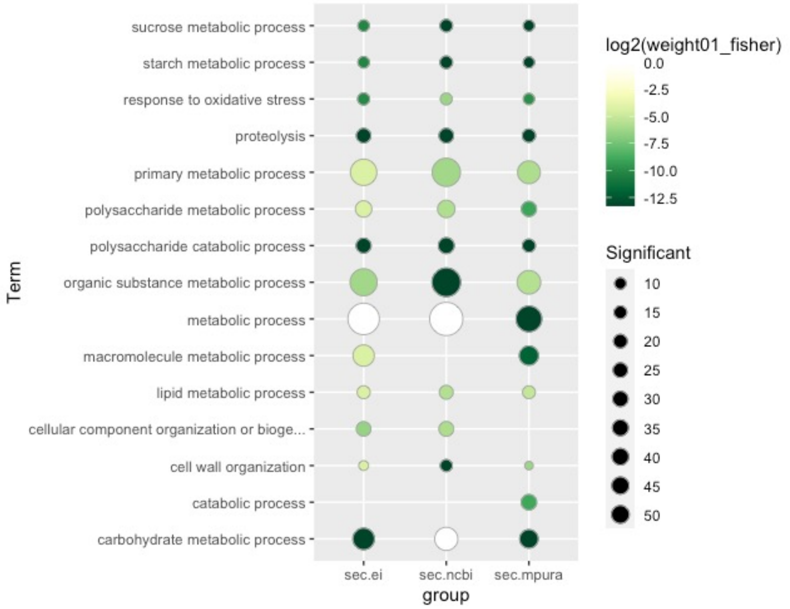
